# High-intensity fear engrams extend beyond threat processing to drive cognitive and affective dysfunction

**DOI:** 10.64898/2026.09.15.751669

**Authors:** Lia P. Iglesias, Fenghua Chen, Claudia R. Cecchi, Maria Luiza T. Novelli, Michel C. van den Oever, Gregers Wegener, Samia Joca

## Abstract

Maladaptive fear memories are a core feature of stress-related disorders and are often accompanied by cognitive and affective disturbances; yet, it remains unclear whether the same neuronal ensembles link maladaptive fear to these behavioural alterations. Here, we show that activity of fear engrams causally contributes to behavioural consequences of intense fear beyond threat processing. Using activity-dependent tagging in the dorsal dentate gyrus (dDG) of mice, we identify neuronal ensembles recruited during high-intensity fear conditioning and show that chemogenetic inhibition of these neurons restores cognitive performance and reduces behavioural despair. These effects are specific to fear-associated ensembles, as inhibition of randomly tagged neurons fails to reproduce them, and depend on persistence of the fear memory, as they are abolished following extinction. Cell-type-specific manipulations revealed that glutamatergic, but not GABAergic, HI-tagged neurons drive fear expression and the associated cognitive and affective alterations, whereas GABAergic neurons exert an opposing influence on fear expression. Consistent with these findings, high-intensity fear was associated with increased reactivation of excitatory fear-tagged neurons and reduced reactivation of GABAergic fear-tagged neurons during behavioural despair. Together, these findings identify high-intensity fear engrams as a causal substrate linking threat-related experiences to cognitive and affective dysfunction and implicate altered excitatory–inhibitory recruitment within a single memory ensemble in the emergence of multi-domain behavioural alterations associated with stress-related disorders.

## Introduction

Conditioned fear is an adaptive process in which neutral cues acquire predictive value for danger, allowing appropriate defensive responses (Gazarini et al., 2023; Trent et al., 2025). These associations are encoded by experience-dependent changes in neuronal activity that promote persistent physical and chemical changes in the brain, engrams (Josselyn et al., 2015; Josselyn & Tonegawa, 2020). Engrams are thought to consist of sparse populations of neurons activated during learning and reactivated during retrieval, which are necessary and sufficient for memory recall (Han et al., 2009; Josselyn & Frankland, 2018; Josselyn et al., 2015; Josselyn & Tonegawa, 2020; Liu et al., 2012). Engram research has provided a mechanistic framework for how pathological fear memories may lead to some of the core symptoms of stress-related mental disorders, such as post-traumatic stress disorder (PTSD), and associated comorbidities, including chronic pain (Stegemann et al., 2023). However, it remains unknown whether fear engrams also drive the cognitive and affective dysfunctions often observed in stress-related mental disorders.

Under conditions such as high-intensity (HI) experiences or prior stress (Gazarini et al., 2023), fear memories generalize across contexts (Cui et al., 2024; Lesuis et al., 2021; Lesuis et al., 2025) and become resistant to extinction (Gazarini et al., 2023). These features are characteristic of PTSD and are thought to underlie its core symptoms, including intrusive responses to neutral cues and avoidance behaviour, as well as their persistence (Lis et al., 2020; Sep et al., 2023). However, PTSD is also associated with cognitive impairments and affective disturbances, frequently meeting criteria for comorbid depression, which is linked to poorer prognosis and reduced treatment response (Flory & Yehuda, 2015). To what extent intense fear memories influence the affective and cognitive domains remain unknown. Previous work has shown that chronic activation of negative memory engrams can induce cognitive deficits through secondary mechanisms involving glial-mediated responses (Jellinger et al., 2024). This interpretation implicitly treats cognitive and potentially affective alterations as downstream consequences of sustained fear-related activity, rather than arising from the acute activity of fear-related neuronal populations themselves.

Here, we tested the hypothesis that the activity of fear engrams mediates cognitive impairment and behavioural despair following HI fear learning. To address this hypothesis, we used an activity-dependent tagging approach in the mice dorsal dentate gyrus (dDG) to identify neurons engaged during fear conditioning and examined their contribution to different fear-induced behavioral outcomes.

## Results

### HI conditioning enhances fear during generalization and after extinction

HI conditioning has been associated with increased fear generalization and reduced extinction (Gazarini et al., 2023). We reason that such fear states may be more likely to impact behavioural domains beyond threat processing, including cognition and affect, whereas low-intensity (LI) fear responses that remain context-specific may not produce these alterations. To examine the impact of conditioning intensity on behavioural and neuroendocrine responses, we compared LI and HI protocols in male and female mice (Fig. 1a). During conditioning, LI and HI groups exhibited freezing responses to foot shocks in both sexes (Fig. 1b,e). However, HI-conditioned males displayed increased freezing across phases, suggesting enhanced fear generalization and reduced extinction (Fig. 1c). These effects were not observed in females (Fig. 1f).

**Figure 1:**
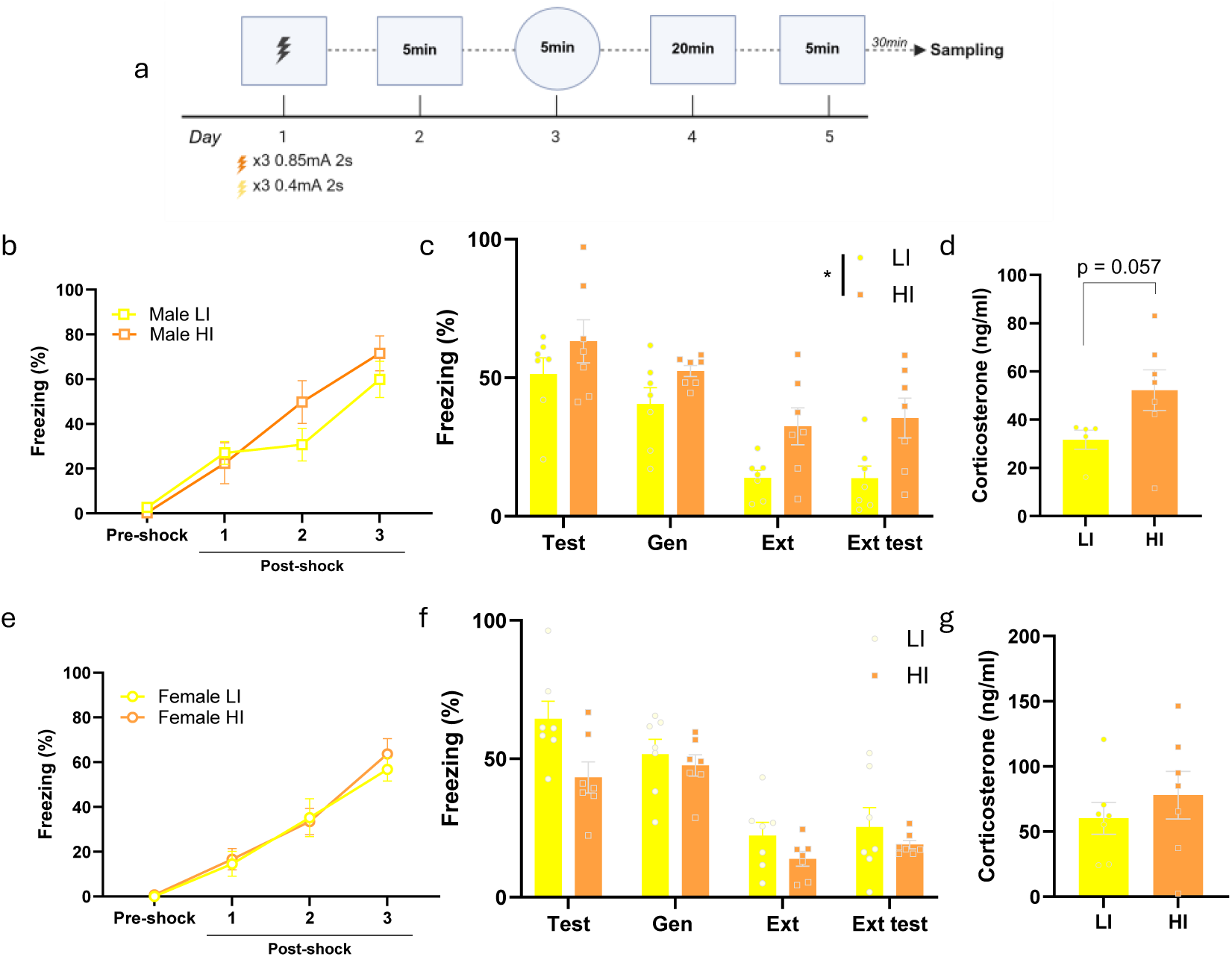
Behavioural and neuroendocrine responses following LI and HI fear conditioning in male and female mice. a Experimental timeline b Freezing levels in males during conditioning. Pre-shock corresponds to the 3 min period before shock delivery; post-shock corresponds to inter-trial intervals, two-way repeated measures ANOVA [Interaction: F(2.796, 33.56) = 1.868, p=0.1571; Time: F(2.796, 33.56) = 42.07, p<0.0001; Intensity: F(1, 12) = 0.8109, p=0.3856], n=7. c Freezing levels in males during test, generalization, extinction, and extinction test sessions, two-way repeated measures ANOVA [Interaction: F(2.267, 27.21) = 0.7884, p=0.4791; Time: F(2.267, 27.21) = 35.94, p<0.0001; Intensity: F(1, 12) = 6.322, p=0.0272], n=7. d Corticosterone levels in males measured 30 min after the extinction test, t-test t-test [t(8.284) = 2.207, p= 0.0572], n=5-7. e Freezing levels in females during conditioning. Pre-shock corresponds to the 3 min period before shock delivery; post-shock corresponds to inter-trial intervals, two-way repeated measures ANOVA [Interaction: F (2.226, 26.72) = 0.2796, p=0.7809; Time: F(2.226, 26.72) = 55.46, p<0.0001; Intensity: F(1, 12) = 0.1732, p=0.6847], n=7. f Freezing levels in females during test, generalization, extinction, and extinction test sessions, two-way repeated measures ANOVA [Interaction: F(2.400, 28.80) = 1.955, p=0.1529; Time: F (2.400, 28.80) = 45.31, p<0.0001; Intensity: F(1, 12) = 3.762, p=0.0763], n=7. g Corticosterone levels in females measured 30 min after the extinction test, t-test [t(10.54)=0.8115, p= 0.4329], n=7. Data are presented as mean ± SEM.

We next assessed whether these behavioural changes were accompanied by sustained physiological responses by measuring corticosterone levels following extinction recall. HI conditioning showed a trend towards increasing corticosterone levels in males, but not females, suggesting persistent fear-related stress-hormone levels despite extinction training (Fig. 1d,g). We next examined components of glucocorticoid signaling associated with stress resilience and vulnerability, including the glucocorticoid receptor (GR) and its co-chaperone FKBP51 (Andero, 2025; Codagnone et al., 2022). In contrast to males, females exposed to HI conditioning showed reduced GR and a trend towards decreased FKBP51 expression (Supplementary Fig. 2a–d), suggesting sex-specific neuroendocrine adaptations.

Together, these results indicate that HI conditioning selectively induces increased fear during generalization and after extinction in males but not in females. Subsequent experiments were therefore restricted to male mice exposed to HI or LI, as they displayed the phenotype needed to compare the neural substrates underlying adaptive versus maladaptive fear memories in our protocol.

### Inhibition of HI-tagged neurons rescues memory impairments and reduces behavioural despair

After fear conditioning with LI or HI, we used an activity-dependent TRAP-based dual-virus system (Matos et al., 2019) to express either mCherry or an inhibitory DREADD (hM4Di-mCherry) in neurons activated during conditioning (Fig. 2a). Inhibition of fear-tagged neurons during retrieval did not reduce freezing (Fig. 2b), suggesting that the tagged population may include functionally heterogeneous neuronal subtypes. In the Novel Object Recognition (NOR), HI-conditioned control animals failed to recognize the novel object, while LI-conditioned animals showed intact performance (Fig. 2c). Inhibition of HI-tagged neurons restored novel object recognition, indicating that activity of this ensemble contributes to memory impairments. The forced swim test (FST) was used to assess changes in behavioural despair of animals exposed to HI and LI. Immobility time did not differ between LI and HI groups in control animals. However, inhibition of HI-tagged neurons reduced immobility time (Fig. 2d), indicating that activity of this ensemble contributes to behavioural despair. These effects were consistent across experimental conditions and were not influenced by test order (Supplementary Fig. 2a-c). Notably, these effects were selective, as inhibition of HI-tagged neurons did not alter social interaction or anxiety-like behaviour (Supplementary Fig. 2d-f).

**Figure 2:**
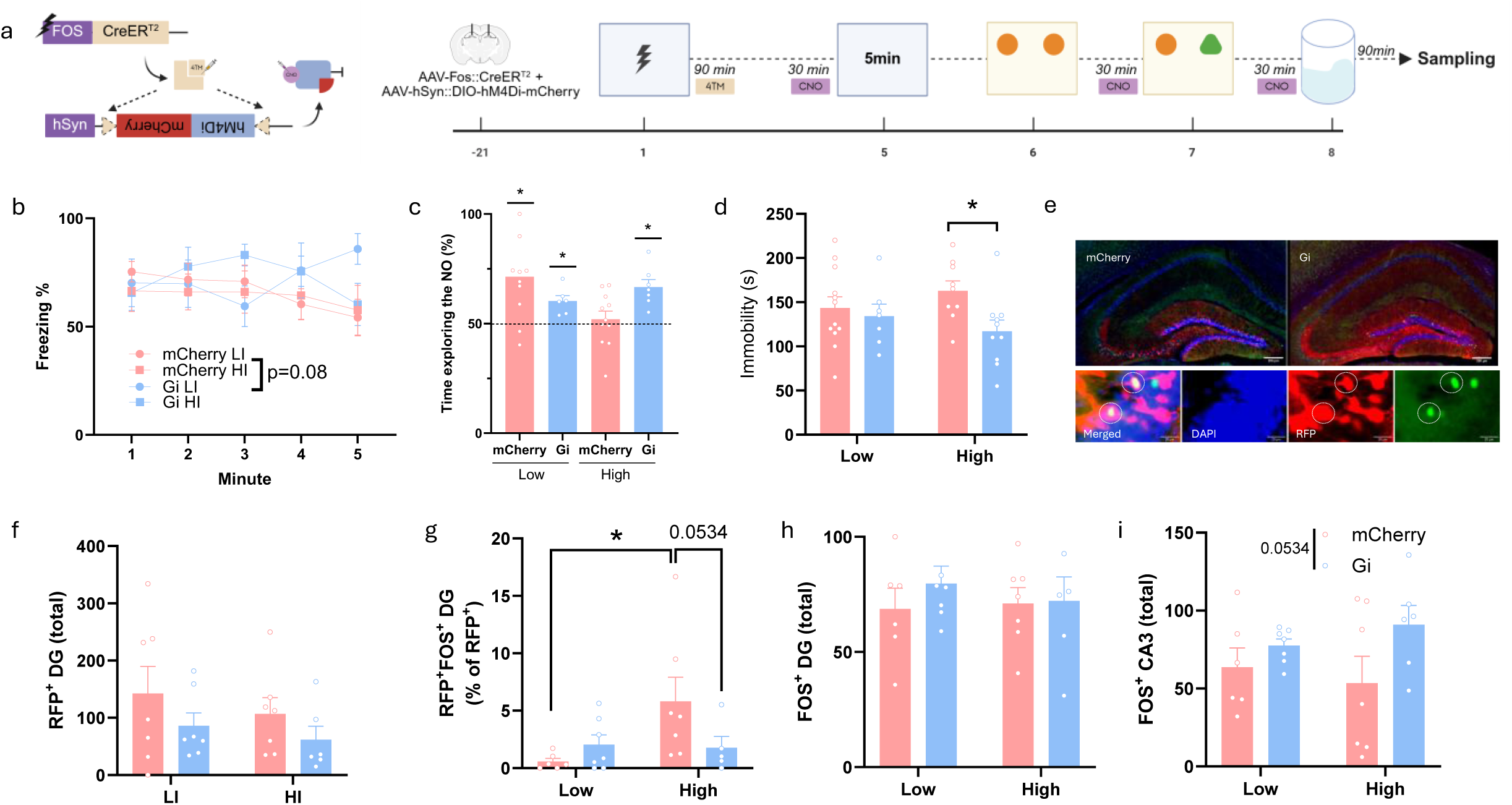
Effects of engram inhibition across behavioural and cellular measures following LI and HI fear conditioning. a Left: schematic of the viral strategy for activity-dependent engram tagging and chemogenetic inhibition. Right: experimental timeline. b Freezing levels by minute during fear recall, three-way mixed effects ANOVA [Time: F(4, 180) = 0.3255, p=0.8606; Intensity: F(1, 180) = 0.07655, p=0.7823; AAV: F(1, 180) = 3.036 p= 0.0831; Time x Intensity: F(4, 180) = 0.775, p=0.5427; Time x AAV: F(4, 180) = 0.8118, p=0.5191; Intensity x AAV: F(1, 180) = 0.1069, p=0.7441; Intensity x AAV x Time: F(4, 180) = 1.410, p=0.2326], n=12-7-11-10. c Percentage of time exploring the novel object in the novel object recognition, one sample t-test [mCherry LI: t(9)=3.404, p=0.0078; Gi LI; t(5)=4.170, p=0.0087; mCherry HI: t(10)=0.5167, p=0.6166; Gi HI: t(6)=4.962, p=0.0025], n=10-6-11-7. d Immobility time in the forced swim test, two-way ANOVA followed by LSD [Interaction: F(1, 34) = 1.948, p=0.1719; Intensity: F(1, 34) = 0.006728, p=0.9351; AAV: F(1, 34) = 4.340, p=0.0448], n=12-7-9-10. e Representative image of engram-tagged neurons (RFP⁺) and Fos⁺ expression. White circles indicate double-positive neurons. Scale bar, 200 µm (HPC) and 20 µm (co-localization). f Number of RFP⁺ cells in DG, two-way ANOVA [Interaction: F(1, 23) = 0.02896, p=0.8664; Intensity: F(1, 23) = 0.8419, p=0.3684; AAV: F(1, 23) = 2.397, p=0.1352], n=7-7-7-6. g Percentage of double-positive (RFP⁺Fos⁺) neurons across experimental groups, two-way ANOVA [Interaction: F(1, 21) = 4.091, p=0.0560; Intensity: F(1, 21) = 3.364, p=0.0808; AAV: F(1, 21) = 0.8916, p=0.3558], n=6-7-7-5. h Number of Fos⁺ cells in the dDG, two-way ANOVA [Interaction: F(1, 22) = 0.3402, p=0.5657; Intensity: F(1, 22) = 0.09307, p=0.7632; AAV: F(1, 22) = 0.5221, p=0.4776], n=6-7-7-6. i Number of Fos⁺ cells in CA3, two-way ANOVA [Interaction: F(1, 22) = 0.8837, p=0.3574; Intensity: F(1, 22) = 0.01595, p=0.9007; AAV: F(1, 22) = 4.166; p=0.0534], n=6-7-7-6. Data are presented as mean ± SEM

Next, we tested the hypothesis that HI conditioning may lead to the formation of less sparse engrams. However, our data show no difference between population size between HI or LI (Fig. 2f). To determine whether the behavioural effects were associated with reactivation of fear-tagged neurons during behavioural despair, we quantified Fos expression in this ensemble following the FST. HI-conditioned animals showed an increased proportion of Fos⁺/RFP⁺ neurons, which was reduced by inhibition of the HI-tagged population (Fig. 2g). Given the role of the DG–CA3 pathway in pattern separation and completion (Lee & Lee, 2020; Lee et al., 2015), we examined Fos expression in these two hippocampal subregions. While the total number of Fos⁺ neurons in the DG was unchanged across groups (Fig. 2h), the inhibition of HI-tagged neurons was associated with a trend towards increased Fos expression in CA3 (Fig. 2i).

Together, these results indicate that activity of HI-tagged neuronal populations contributes to cognitive and affective alterations following HI fear conditioning.

### Random neuronal inhibition does not reproduce the effects of HI-tagged neuron inhibition

To test whether the behavioural rescue was specific to inhibition of HI-tagged neurons, we tagged neurons activated in a neutral context (Fig. 3a) using the same activity-dependent viral strategy (Fig. 2a). Animals were subsequently subjected to HI fear conditioning, and random-tagged neurons were inhibited during fear retrieval, NOR, and FST. Inhibition of randomly tagged neurons did not alter freezing during fear retrieval (Fig. 3b). In the NOR task, inhibition did not restore novel object discrimination (Fig. 3c), indicating that non-specific neuronal inhibition is insufficient to rescue fear-induced memory impairments. In the FST, inhibition of randomly tagged neurons increased immobility time (Fig. 3d). The size of the random engram does not significantly differ from the HI fear-tagged neuronal population (Fig. 3f)

**Figure 3:**
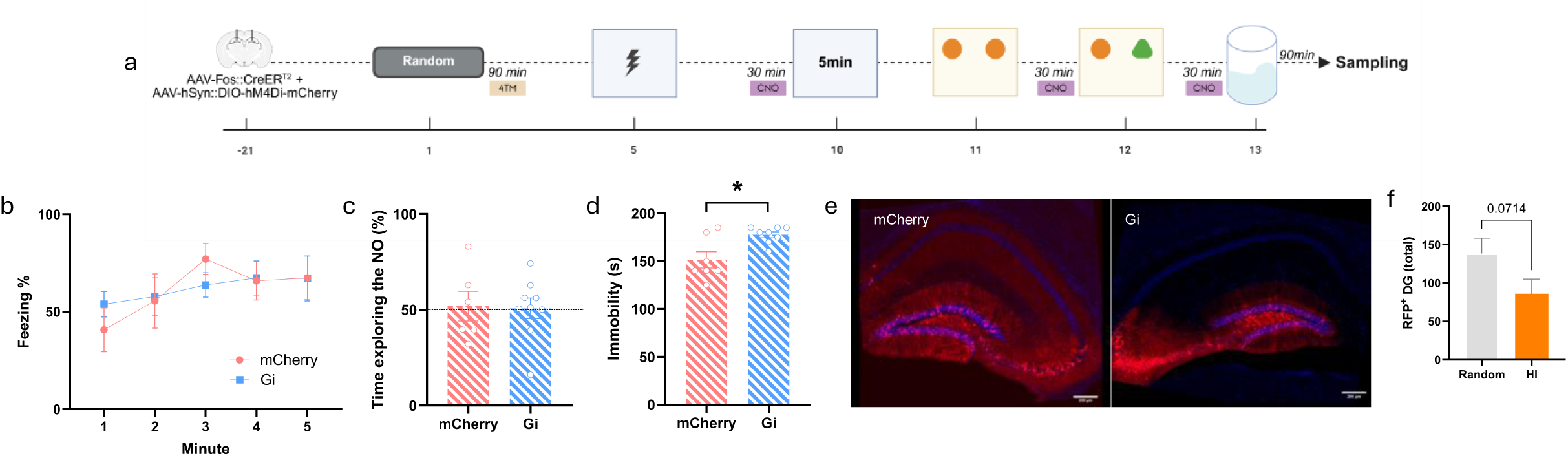
Effects of random engram inhibition across behavioural tasks. a Experimental timeline. b Freezing levels by minute during fear recall, two-way repeated measures ANOVA [Interaction: F(3.135, 43.89) = 0.5025, p=0.6905; Time: F(3.135, 43.89) = 2.051, p=0.1181 ; AAV: F(1, 14) = 0.008433, p=0.9281], n=7-9. c Percentage of time exploring the novel object in the novel object recognition, one sample t-test [mCherry: t(5)=0.2437, p=0.8172; Gi: t(8)=0.1363, p=0.8949], n=6-9. d Immobility time in the forced swim test, t-test [t(13)=3.033, p=0.0096], n=7-8. e Representative image of engram-tagged neurons (RFP⁺). Scale bar, 200µm. f Number of RFP⁺ cells in DG, t-test [t(27)=1.876, p=0.0714], n=17-13. Data are presented as mean ± SEM.

Together, these findings indicate selective inhibition of the HI-tagged population in the dDG rescues cognitive and affective alterations associated with HI.

### Inhibition of HI-tagged neurons improves discrimination index

To determine whether the HI neuronal population involved in cognitive impairments and behavioural despair is also involved in inappropriate fear responses, neurons activated during HI conditioning were tagged using the same activity-dependent viral strategy (Fig. 2a) and inhibited during fear retrieval and generalization (Fig. 4a). Our data shows that inhibition of HI-tagged neurons reduced freezing during the generalization test and improved context discrimination (Fig. 4c,d).

**Figure 4:**
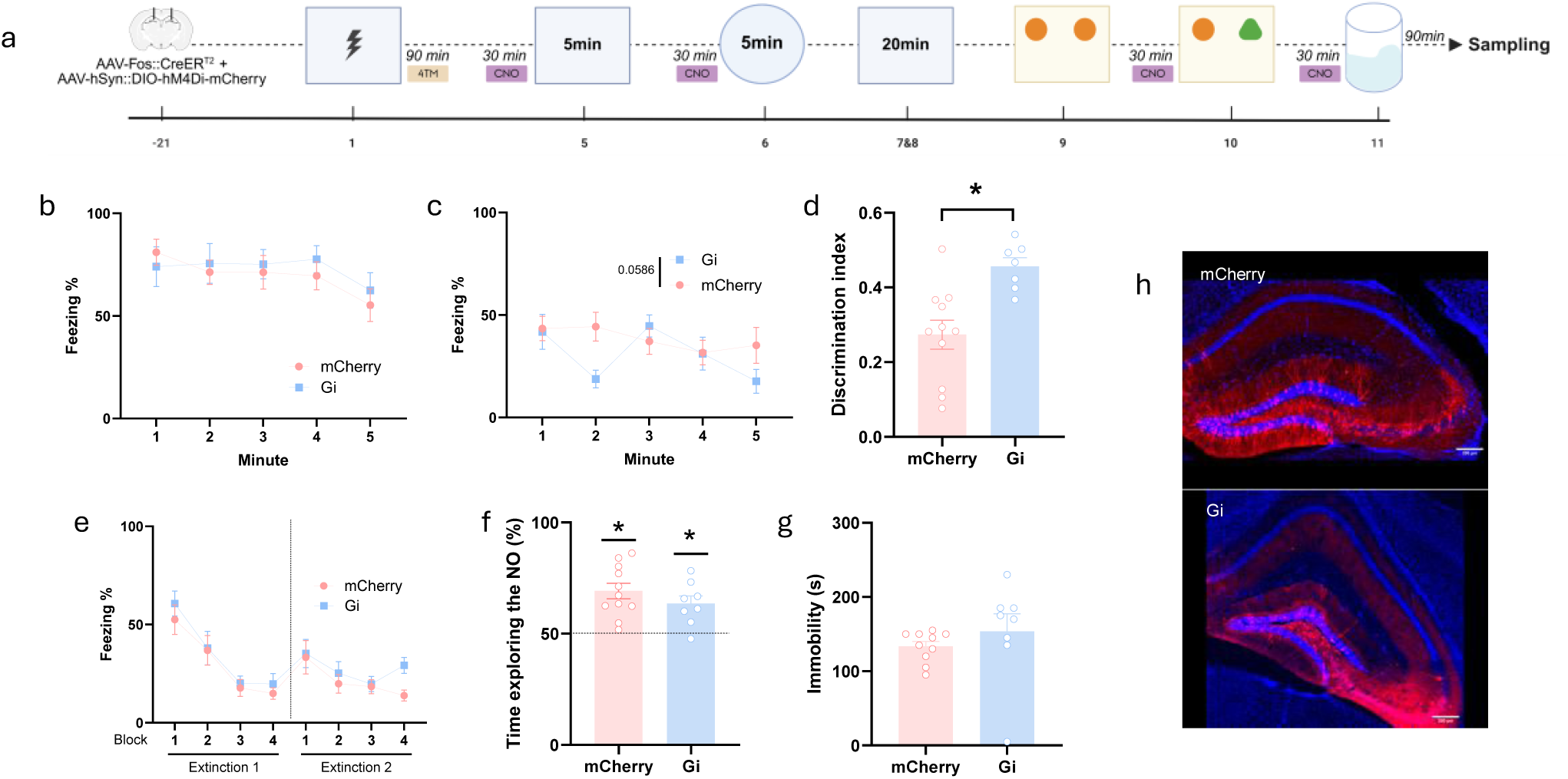
Effects of extinction in engram inhibition across behavioural tasks. a Experimental timeline. b Freezing levels by minute during fear recall, two-way repeated measures ANOVA [Interaction: F(3.003, 51.06) = 0.4882, p=0.6922; Time: F(3.003, 51.06) = 2.747, p=0.0523; AAV: F(1, 17) = 0.1761, p=0.6800], n=11-8. C Freezing levels by minute during fear generalization, two-way repeated measures [Interaction: F(3.229, 54.89) = 1.759, p=0.1621 ; Time F(3.229, 54.89) = 1.821 p=0.1501; AAV: F(1, 17) = 4.111, p=0.0586], n=11-8. d Discrimination index, t-test [t(16)=3.471, p=0.032], n=11-7. e Freezing levels by block (5min) during two sessions of fear extinction, two-way repeated measures ANOVA [Interaction: F(7, 119) = 0.4319’, p=0.8805; Block: F(7, 119) = 13.81, p<0.001; AAV: F(1, 17) = 1.189, p=0.2908], n=11-8. f Percentage of time exploring the novel object in the novel object recognition, one sample t-test [mCherry: t(10)=5.507, p=0.0003; Gi: t(7)=3.924, p=0.0057], n=11-8. g Immobility time in the forced swim test, t-test [t(16)=0.9135, p=0.3745], n=10-8. h Representative image of engram-tagged neurons (RFP⁺). Scale bar, 200µm. Data are presented as mean ± SEM.

### Extinction abolishes the behavioural effects of HI-tagged neuronal inhibition

To assess whether the cognitive and affective alterations depend on the persistence of the fear memory, we tested whether successful fear extinction would blunt the behavioral effects induced by inhibition of HI-tagged neurons. After generalization, animals underwent extinction training. After extinction, tagged neurons were inhibited during NOR and FST. Extinction learning was not affected by prior inhibition, as freezing levels during extinction training were comparable between groups (Fig. 4e). Following extinction, animals showed intact object recognition, and inhibition of HI-tagged neurons did not alter performance in the NOR or immobility time in the FST (Fig. 4f,g). These results indicate that the cognitive and affective deficits associated with HI fear depend on the persistent activity of the fear engram.

### Cell-type-specific inhibition reveals opposing roles of glutamatergic and GABAergic HI-tagged neurons

Inhibition of HI-tagged neurons rescued memory impairments and reduced behavioural despair, while paradoxically not decreasing freezing during fear retrieval (Fig. 2). Given that engram activation is required for memory expression (Josselyn & Frankland, 2018; Josselyn et al., 2015; Josselyn & Tonegawa, 2020), these findings raise the possibility that the tagged population may comprise functionally distinct neuronal subtypes with opposing roles. To test that, we combined TRAP-induced activity-dependent tagging with cell-type-specific targeting strategies and selectively manipulated glutamatergic or GABAergic neurons activated during HI conditioning (Fig. 5a). Glutamatergic neurons were targeted using a CaMKIIα-driven construct, whereas GABAergic neurons were targeted using a Dlx-based approach.

**Figure 5:**
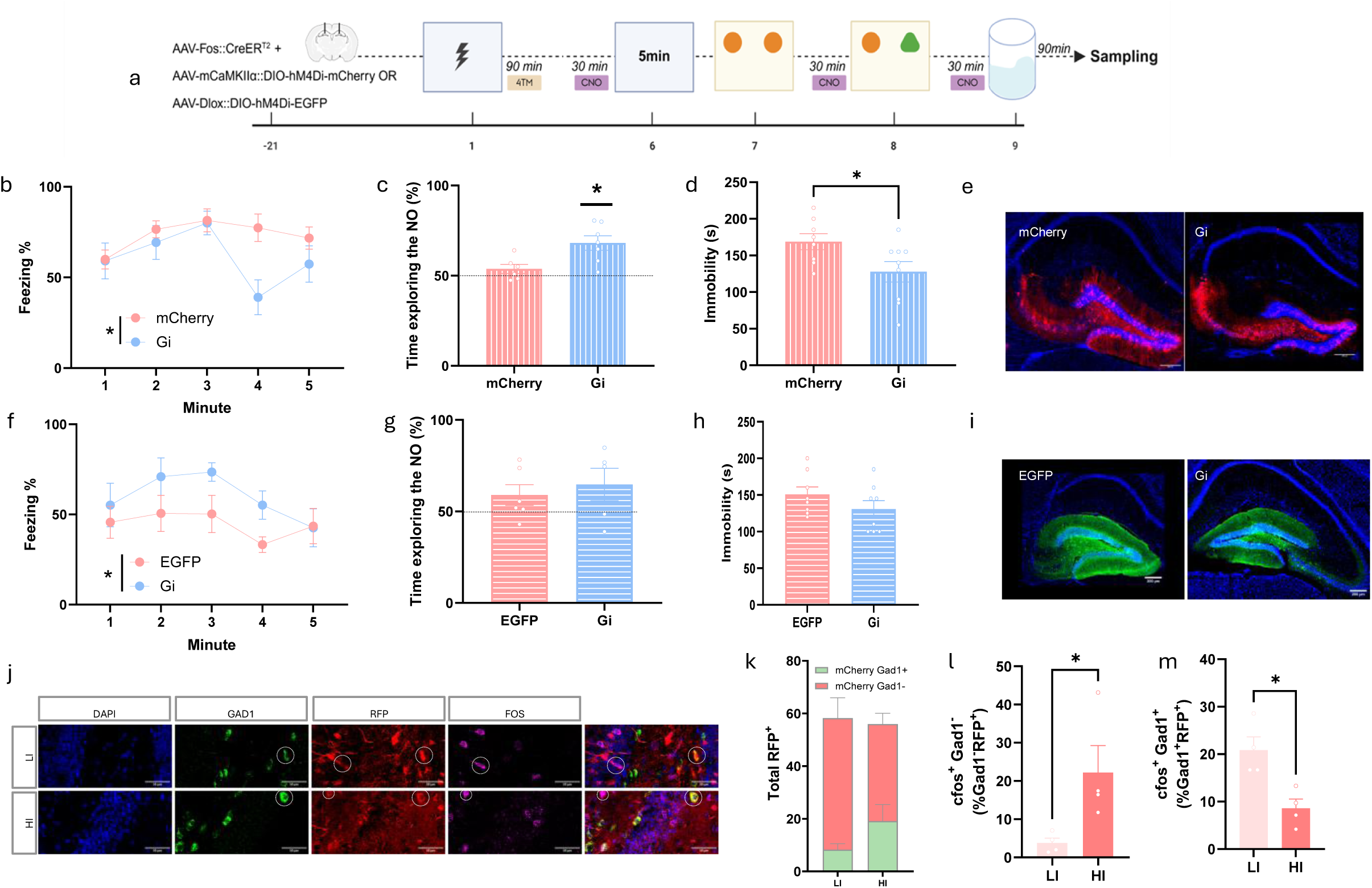
Effects of the inhibition of glutamatergic and gabaergic neurons active during fear across behavioural measures. a Experimental timeline. b Freezing levels by minute during fear recall, two-way repeated measures ANOVA [Interaction: F(4, 70) = 1.743, p=0.1503; Time: F(4, 70) = 2.644, p=0.0406; AAV: F(1, 70) = 5.646, p=0.0202], n=8-9. c Percentage of time exploring the novel object in the novel object recognition, one sample t-test [mCherry: t(5)=1.478, p=0.1994 ; Gi: t(6)=4.465, p=0.0043], n=6-7. d Immobility time in the forced swim test, t-test [t(15)=2.257, p=0.0394], n=8-9. e Representative image of engram-tagged neurons (RFP⁺). Scale bar, 200µm. f Freezing levels by minute during fear recall, two-way repeated measures ANOVA [Interaction: F(4, 65) = 0.6289, p=0.6436; Time F(4, 65) = 1.865, p=0.1273; AAV: F(1, 65) = 6.447, p=0.0135], n=8-7. g Percentage of time exploring the novel object in the novel object recognition, one sample t-test [EGFP: t(5)=1.573, p=0.1765; Gi t(4)=1.661, p=0.1721], n=6-5. h Immobility time in the forced swim test, t-test [t(14)=1.296, p=0.2161], n=8. i Representative image of engram-tagged neurons (EGFP⁺). Scale bar, 200µm. j Representative image of engram-tagged neurons (RFP⁺), Gad1⁺ and cfos⁺ expression. White circles indicate double-positive neurons. Scale bar, 50 µm. k Total number of RFP⁺ and Gad⁺ RFP⁺ neurons, two-way ANOVA [Interaction: F(1, 12) = 4.765, p=0.0496, Intensity: F(1, 12) = 0.04188, p=0.8413; Cell type: F(1, 12) = 29.29, p=0.0002], n=4-4. l Percentage of cfos+gad1-over gad1-RFP+ neurons, t-test [t(6)=2.558, p=0.043], n=4-4. m Percentage of cfos+gad1+ over gad1+RFP+ neurons, t-test [t(6)=3.579, p=0.0117], n=4-4. Data are presented as mean ± SEM.

Inhibition of HI-tagged glutamatergic neurons reduced freezing during fear retrieval, restored performance in the NOR task and reduced immobility time in the FST (Fig. 5b-d). In contrast, inhibition of HI-tagged GABAergic neurons increased freezing during fear retrieval (Fig. 5e), without affecting performance in the NOR or FST (Fig. 5f,g), suggesting that they restrain fear expression without contributing to its associated cognitive and affective deficits.

To further investigate how the gabaergic neurons were engaged during the behavioural alterations induced by HI fear, we performed an RNAscope (Fig. 5j) in animals conditioned with LI or HI (Fig. 2). We observed again that there is no difference in the size of the tagged populations between intensities. Similarly, we found no statistically significant difference in the number of tagged neurons expressing the gabaergic marker gad1 (Fig. 5k). Interestingly, while a higher number of non-gabaergic fear-tagged neurons were active during the FST in HI-exposed animals (Fig. 5l), fear-tagged neurons co-expressing gad1 were hyp0active in the HI compared to LI (Fig. 5m).

Together, these results indicate that glutamatergic HI-tagged neurons constitute the core of the fear engram, with their activity driving both fear expression and the associated cognitive and affective alterations. GABAergic neurons recruited during fear, in contrast, exert an opposing influence on fear expression and remain hypoactive during cognitive and affective-related behaviours.

## Discussion

Maladaptive fear memories are a central feature of stress-related disorders and are frequently accompanied by cognitive and affective alterations, which are often considered secondary consequences of the disorder rather than directly linked to the dysregulation of the fear memory itself. Here, we show that HI fear conditioning enhanced fear responses during generalization and after extinction and recruits hippocampal neuronal populations whose activity contributes to memory impairments and behavioural despair. Accordingly, inhibition of hippocampal neurons tagged during HI fear learning restored cognitive performance and reduced behavioural despair. These effects were specific to such fear-associated neuronal populations and depended on the persistence of the fear memory, as they were abolished following extinction. Furthermore, cell-type-specific manipulations of the tagged neuronal population revealed that glutamatergic, but not GABAergic neurons, constitute the fear engram, which is also the neural substrate mediating the behavioural impairments associated with fear exposure. Finally, the gabaergic neurons recruited during HI conditioning presented lower activity during behavioural despair in comparison with the same population in the LI group and with the glutamatergic counterpart. Together, these findings support a model in which activity of HI fear engrams and hypoactivity of its inhibitory component links threat-related experiences to its associated cognitive and affective dysfunction.

Testing how adaptative versus maladaptive memories relate to fear-induced behavioral consequences first required a behavioural approach able to distinguish between context-specific, extinction-sensitive fear responses and generalized, extinction-resistant fear, which we achieved by varying the intensity of the aversive stimulus (Cui et al., 2024; Dos Santos Correa et al., 2019; Gazarini et al., 2023). Consistent with previous reports (Andero, 2025; Day & Stevenson, 2020), HI conditioning induced fear generalization and resistance to extinction in males, but not in females tested under the same protocol. These behavioural differences parallel distinct neuroendocrine profiles, whereby females show reduced FKBP51 and glucocorticoid receptor expression, a pattern often associated with increased resilience to stress (Andero, 2025; Codagnone et al., 2022). These findings highlight sex-specific adaptations that possibly moderate female vulnerability to HI fear in our protocol. Contrasting results regarding sex-vulnerability in rodent models of fear learning, however, implicate more complex mechanisms depending on the paradigm used (Binette et al., 2022; Day & Stevenson, 2020; Riccardi et al., 2024). Since our main goal was test whether maladaptive fear memories triggered by intense fear would underlie the development of behavioral deficits, rather than sex-driven vulnerability, we focused our subsequent experiments on male mice, as they reliably displayed the fear generalization and extinction-resistance phenotype required for this comparison.

We next investigated whether neuronal populations recruited during HI fear influence behavioural domains beyond threat processing. We focused on the dDG, a region implicated in fear generalization (Cui et al., 2024; Jeong et al., 2024; Lesuis et al., 2021), extinction (Bernier et al., 2017; Gong et al., 2022; Lacagnina et al., 2019), recognition memory (Castillon et al., 2026; Hainmueller & Bartos, 2020; van Dijk & Fenton, 2018) and affective disturbances (Sun et al., 2023; Umschweif et al., 2021). Although the role of dDG ensembles in fear retrieval remains debated (Carretero-Guillén et al., 2024; Denny et al., 2014; Madroñal et al., 2016), we expected that inhibition of fear-tagged neurons would impair fear retrieval regardless of conditioning intensity, confirming their participation in the fear engram (Carretero-Guillén et al., 2024; Josselyn & Frankland, 2018; Josselyn et al., 2015; Josselyn & Tonegawa, 2020). However, this was not observed, suggesting that the tagged population comprises functionally heterogeneous neuronal ensembles. Notably, the HI-tagged neuronal populations were overactive during the FST and their selective inhibition attenuated the behavioural consequences induced by intense fear in both the FST and NOR. Therefore, whereas prior work has shown that fear-associated cognitive deficits arise only after long-term activation of negative-memory engrams, potentially through glial-dependent mechanisms (Jellinger et al., 2024), our results indicate that the behavioral deficits arise directly from spontaneous overactivity of fear-associated neuronal populations. Our data further indicates that the behavioural alterations associated with HI fear cannot be attributed to the recruitment of an enlarged neuronal population, as no differences in ensemble size were observed between the LI and HI groups. This finding is consistent with previous studies showing that variations in fear intensity are not accompanied by changes in DG engram size (Cui et al., 2024).

Inhibition of fear-tagged neurons resulted in increased Fos expression in CA3, without significant changes in the DG. The lack of significant changes in the DG Fos is not necessarily surprising if we consider that only a small number of neurons is often tagged by fear, while a much larger untagged granule cell population can still be free to fire normally, thereby preventing bulk DG Fos counts to reach significance. Nevertheless, the inhibition of HI-tagged cells in the DG resulted in disinhibition of neuronal populations within CA3. DG–CA3 pathway is key in pattern separation and memory processing (Lee & Lee, 2020; Lee et al., 2015), with CA3 activity increasing when a context is recognized as novel (Hainmueller & Bartos, 2018). These findings raise the possibility that activity of fear-associated neuronal ensembles in the DG modulates downstream hippocampal processing, but further studies are required to understand how this can relate to our behavioral findings given the complexity of DG-CA3 circuitry (Basu & Siegelbaum, 2015).

Notably, our experiments using inhibition of randomly tagged neurons failed to reproduce the effects induced by inhibition of fear-tagged neurons, indicating that the behavioural deficits arise specifically from the fear-associated neuronal populations rather than from a generic mnemonic disruption or non-specific silencing of fear-related activity. In fact, non-specific neuronal inhibition did not restore recognition memory and increased behavioural despair, consistent with a broader role of DG activity in behavioral adaptation to stress (Gergues et al., 2021; Tunc-Ozcan et al., 2019).

In addition, if cognitive and affective alterations are merely driven by sustained fear, then reducing fear through extinction learning should naturally decrease the activity of HI-tagged neurons and abolish behavioural alterations (Lacagnina et al., 2019; Luft et al., 2024). Our findings demonstrate that extinction abolished both cognitive impairments and the impact of HI-tagged neuron inhibition, indicating that these alterations are maintained by ongoing engram activity rather than by downstream processes that become independent of the original memory trace.

Finally, we addressed the observation that inhibition of HI-tagged neuronal ensembles did not impair retrieval. Neuronal populations activated during learning are not homogeneous but instead comprise distinct subpopulations that may encode different features of the experience or serve different functional roles (Pouget et al., 2026). In particular, excitatory principal neurons are widely considered to constitute the core component of the memory engram (Josselyn et al., 2015; Josselyn & Tonegawa, 2020; Lesuis et al., 2025), whereas inhibitory neurons recruited during learning regulate engram formation, sparsity, specificity and retrievability (Lesuis et al., 2025; Tome et al., 2024; Wu et al., 2026). We selectively targeted glutamatergic and GABAergic neurons activated during HI conditioning. Inhibition of glutamatergic HI-tagged neurons reduced freezing during retrieval, consistent with their contribution to the fear engram, and prevented memory impairments and reduced behavioural despair. In contrast, inhibition of GABAergic HI-tagged neurons increased freezing without affecting cognitive performance or behavioural despair. It is possible that altered balance between excitatory and inhibitory components within these ensembles influences the extent to which engram activity generalizes across behavioural domains. Indeed, we observed a marked reduction in reactivation of GABAergic fear-tagged neurons together with increased reactivation of non-GABAergic fear-tagged neurons during the FST. Together with the opposing effects of cell-type-specific inhibition, these findings support a model in which maladaptive fear arises from an altered balance between inhibitory and excitatory components of the fear-recruited neuronal population, promoting inappropriate engagement of the fear engram outside threat-related contexts.

In summary, we demonstrate that DG engrams encoding HI fear mediate cognitive and affective alterations beyond threat processing. Our data provides a new mechanism for how intense fear causes cognitive and affective impairments, bringing new insights into the neurobiology of stress-related disorders and its comorbidities.

## Methods

### Animals

Male and female, 8-weeks-old C57BL/6RccHsd (Envigo) were housed grouped in Eurostandard Type IIIH cages with raised lids, shelter, and nesting material, with free access to food and to tap water. The cages were kept in a temperature-controlled room (21±1°C) with a standard dark-light cycle of 12/12h (lights on 06:00), and all experiments were performed during the light phase of the cycle. All procedures performed were approved by the Danish National Committee for Animal Experimentation (2023-15-0201-01523) in agreement with the EU Directive 2010/63/EU and the Danish Law (LBK nr 1107 af 01/07/2022).

### AAV and stereotaxic surgery

AAV5-Fos::CreERT2 (gift from Van den Oever M., titer: 1.2 × 10^13^) and Cre-dependent AAVs (Zurich Viral Facility, titers: 5.0–6.0 × 10^12^) encoding designer receptor exclusively activated by designer drugs (DREADD) fused to a fluorescent reporter AAV5-hSyn::DIO-hM4Di-mCherry, AAV5-CaMKIIα::DIO-hM4Di-mCherry, AAV5-dlox::DIO-hM4Di-EGFP or a fluorescent reporter AAV5-hSyn::DIO-mCherry, AAV5-CaMKIIα::DIO-mCherry, AAV5-dlox::DIO-EGFP were infused as previously described (Matos et al., 2019). Briefly, animals were anesthetized with inhalatory sevoflurane and local lidocaine (2%) and placed in the stereotaxic apparatus (RWD71000). A virus mixture (0.5μL/hemisphere) of AAV-Fos::CreERT2 and Cre-dependent AAV (ratio 1:500; AAV-Fos::CreERT2 final titer of 2.4 × 10^10^) was injected into the dorsal HPC (AP+1.9, ML ±1.6, DV-2 from lambda) at a flow rate of 0.1μL/min, the glass pipette was removed 5min after the infusion was finished. Behavioural experiments were conducted 21 days after surgery.

### Drugs

Treatments were done as previously described (Matos et al., 2019). 4-hydroxytamoxifen (4TM, Sigma Aldrich) was diluted in 5% DMSO and 1% Tween80 in saline. 4TM (25 mg/kg) was injected 2h after fear conditioning to allow targeted recombination followed by a quarantine period of 5 days. Clozapine N-oxide (CNO, Tocris) was diluted in saline. CNO (5 mg/kg) was injected 30 min before behavioural test to activate the inhibitory DREADD. 4TM and CNO were injected at 10ml/kg i.p.

### Contextual Fear Conditioning

Mice underwent contextual fear conditioning in a squared box with stainless-steel grid floor inside a soundproof cabinet (Multi Conditioning System, Version 1.0, TSE Systems GmbH, Bad-Homburg, Germany) with constant noise (50-60dB) and control light (200lx). Mice were placed in the conditioning box and let undisturbed for 120s, after which they received a series of three footshocks of 2 s (LI: 0.45mA; HI: 0.85 mA) separated apart for 40 and 30s, animals were returned to their home-cage 60s after the last footshock. Test was performed by exposing the animals to the squared conditioning context for 5 min in the absence of footshocks. Extinction sessions consist of one or two 20 min exposure to the squared conditioning context in the absence of footshocks. Generalization test was performed by exposing the animals to a transparent cylinder with black floor. Freezing behaviour was analyzed by TSE Systems multiconditioning software.

### Novel Object Recognition (NOR)

During training animals were placed in a grey squared context with two identical objects, animals were removed from the arena when reached 30 s of exploration (Lueptow, 2017). Exploration was considered when the animal was directly interacting with the object or with the nose in the direction of the object (1 cm); climbing the object or touching it to explore above it was not considered exploration time. The test was performed 24h after the training session. During test, the animal was placed in the same context with one of the objects from the training session and a new object, animals were removed from the arena after 10min. The session was recorded and the first 30 s of exploration were manually analyzed by a blinded experimenter (Lueptow, 2017). Animals that failed to reach 30s of exploration in 10 min in either of the steps were excluded (Lueptow, 2017).

### Forced Swimming Test (FST)

Animals were placed in a transparent cylinder of 18 cm of heigh filled with 24±0.5°C water (13.5 cm) for 5 min. To eliminate olfactory cues, the water was changed between each session. The session was recorded on video and immobility, and swimming were manually evaluated by a blinded experimenter (Silva et al., 2025).

### Random tagging

Animals were exposed to a grey rectangular context (40x10x25 cm, 100 lux) for 5 min. Social interaction test

The animals were placed for 2 min in an empty arena (50x50x40 cm). After 2 min a conspecific of the same sex and matching dimension and age was placed in a perforated box in the arena. The animal was allowed to interact with the conspecific for 5 min. The session was recorded and the time interacting was manually analyzed by a blinded experimenter (Berton et al., 2006).

### Elevated plus maze

The apparatus consisted of a plus-shaped maze (25x15x20 cm) elevated above the floor (30 cm), with two opposite open arms and two opposite closed arms connected by a central platform. At the beginning of the test, each mouse was placed in the center of the maze facing an open arm and allowed to freely explore the apparatus for 5 min. Sessions were recorded from the top and time in the open and close arms as well as entries in the arms were manually quantified by a blinded experimenter.

### Perfusion and tissue preparation

Transcardial perfusion was performed 30 min (RNAscope) or 90 min (immunofluorescence) after behaviour. Animals were perfused with 50ml of ice-cold PBS at 10ml/min flowrate. Brains were extracted and post-fixed with PFA 4% for 48 h followed by 48 h in sucrose 30%. Brains were frozen in isopentane. For RNA scope brains were kept at -80°C, cut at 15μm in a cryostat and directly mount into slides. For immunofluorescence, brains were kept at -20°C, cut at 50μm in a cryostat and kept in cryoprotective solution.

### Immunofluorescence

Sections (n=5-7/animal) were permeabilized in 1 % Triton X-100 in PBS (PBS-T) for 15 min, 0.75% glycine for 10 min and washed in PBS twice for 5 min. Then, sections were blocked for 60 min with 5% Bovine Serum Albumin (BSA) in PBS-T and incubated overnight (4°C) with RFP (1:1000; Rockland 600-401-379) and cFos (1:500; SYSY No226017) in 5% BSA in PBS. After extensive washing, sections were incubated with Alex Fluor 488 goat anti-rat IgG (1:500; Invitrogen, A11006) and Alexa red 555 anti-rabbit (1:1000; abcam, ab150078) for 2hrs at room temperature. Finally, the sections were washed in PBST (3 × 10 min) and incubated with Dapi (Sigma aldrich).

### RNAscope

RNAscope fluorescent in situ hybridization was performed using the RNAscope Multiplex Fluorescent Reagent Kit v2 (Bio-Techne) according to the manufacturer’s instructions. Briefly, brain sections were processed using the Pretreat-Pro protocol and hybridized with probes targeting Gad1 (NM_008077, cat. 316921) and cfos (NM_010234, cat. 400951), 1:50. Probes were visualized using ClariTSA™ fluorophore 520 (1:1000) and 650 (1:3000) respectively. Following RNAscope labeling, sections were subjected to immunofluorescence staining using the antibodies described above at the same concentrations and incubation times.

### Imaging

For immunofluorescence imaging was done using an Olympus SLIDEVIEW VS200 slide scanner at x20 or x40 magnification. A 50 μm z-stack combined image was obtained from every slice. Quantification of immunopositive cells per HPC field was done semi-automatically with Qupath (Courtney et al., 2022). Double-positive cells were manually counted by an investigator blinded to experimental condition. For RNAscope imaging was done using the Andor spinning disk confocal and the software FUSION at x20 magnification. A 0.5 µm z-stack combined image was obtained from every slice. Quantification of positive cells was performed manually in QuPath by an experimenter blinded to the experimental conditions.

### Quantification of Corticosterone levels by ELISA

Blood samples were collected by cardiac puncture 30 min after behaviour. The plasma samples were stored in -80°C. Corticosterone levels were measured using a commercial ELISA kit (ADI-900-097, ENZO Life Science, NY, USA) according to the manufacturer’s protocol for small volumes. The samples were diluted 1:40 in steroid displacement reagent (SDR 1:100 in ultrapure water). The optical density was measured using a Clariostar microplate reader (BMG LabTech, Germany) at 405 nm and the results were calculated using a four-parameter logistic curve fit.

### Western Blot

The HPC was dissected 30 min after behaviour, weighed and homogenized in ice-cold RIPA buffer (50 mM Tris-HCl pH 7.4, 150 mM NaCl, 1% NP-40, 0.1% Sodium Dodecyl Sulfate - SDS, 0.5% Sodium deoxycholate, 10% Glycerol) 20:1 (w/v), supplemented with 1mM NaF 1mM, 2mM Na3VO4 and 1% HALT protease 1%. Tissue homogenization was performed using the Precellys Evolution homogenizer (two cycles of 15 seconds at 5000 rpm), followed by 15 min centrifugation at 13.000 rpm. Total protein was determined using the commercial BCA assay kit (Pierce BCA protein assay kit, Thermo Scientific) at 562 nm absorbance. The samples were kept at -80°C.

Cell lysates were denatured at 65°C for 15min in sample buffer (3M Tris-HCl ph6.8, 10% SDS, 100% Glycerol, 2,5% Bromophenol blue and 1.25M Dithiothreitol - DTT in ultrapure water) and 30μg of protein was loaded per well on a NuPAGE 10% Bis-Tris Midi Gel (Invitrogen). Gel electrophoresis was performed in MOPS running buffer (Invitrogen) using PowerEase™ Touch 350W Power Supply (Thermo Scientific) at 100V for 60 min, transferred to a 0.2um nitrocellulose membrane (Trans-Blot Turbo Transfer Pack, Bio-Rad) using TransBlot Turbo Transfer System (Bio-Rad) and blocked in Intercept Blocking Buffer (IBB) (LI-COR Bioscience). Primary Antibodies including glucocorticoid receptor mouse monoclonal antibody (1:500; #47411, Cell Signaling) 1:500, FKBp51 (1:1000; ab126715, Abcam) 1:1000 and β-actin mouse monoclonal antibody (1:3000; 926-42212, LI-COR Biosciences) 1:3000 were diluted in IBB 0.1% Tween 20, added to the membranes and incubated at 4°C overnight, followed by incubation with IRDye 680RD goat anti-mouse or 800CW goat anti-rabbit IgG secondary antibodies (1:20000 in IBB/0,1% Tween20) for 1 h, at room temperature. The images were acquired using an infrared laser system (Odyssey CLx Image, LI-COR Bioscience) and the protein was quantified using the image studio software version 5.0.

### Statistical Analysis

Statistical analyses were performed using GraphPad Prism. Normality was assessed using Shapiro-Wilk test, and outliers were identified using Grubbs’ test. Comparisons between two groups were performed using unpaired Student’s t-tests. The NOR data was evaluated by comparison against a theoretical null value using one-sample t-tests. For experiments involving more than two groups or multiple factors, one-way ANOVA, two-way ANOVA, or repeated-measures ANOVA was used. When significant main effects or interactions were detected, appropriate multiple-comparison post hoc tests were performed to identify group differences. Data are presented as mean ± SEM, statistical significance was set at p < 0.05.

## Supporting information

Supplementary results

## Acknowledgements

This work was supported by the European Union under the Marie Curie Postdoctoral fellowship (grant number HORIZON-MSCA-2022-PF-01-01, ID:101110693 to ILP) and AP Muller Fonden (to SJ and ILP).

