## Supplementary results for "High-intensity fear engrams extend beyond threat processing to drive cognitive and affective dysfunction"

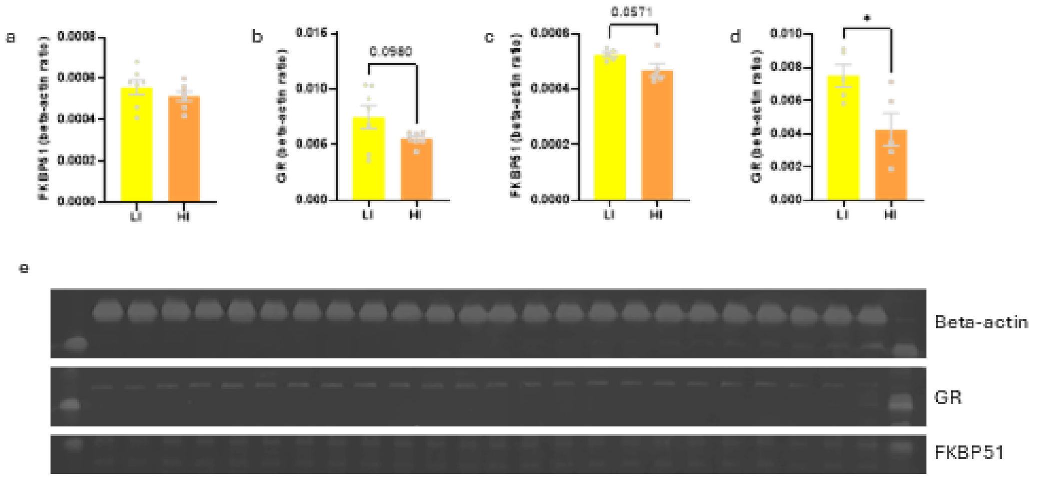


Supplementary figure 1: Neuroendocrine targets in males and females conditioned with LI or HI. a FKBP51 levels in the hippocampus of males measured 30 min after the extinction test, t-test [t(12)=1.024, p=0.3259], n=7. b GR levels in the hippocampus of males measured 30 min after the extinction test, t-test [t(12)=1.794, p=0.0980], n=7. c FKBP51 levels in the hippocampus of females measured 30 min after the extinction test, t-test [t(8)=2.221, p=0.0571], n=5. d GR levels in the hippocampus of females measured 30 min after the extinction test, t-test [t(8)=2.734, p=0.0257], n=5. e Western Blot membranes. Data are presented as mean ± SEM.

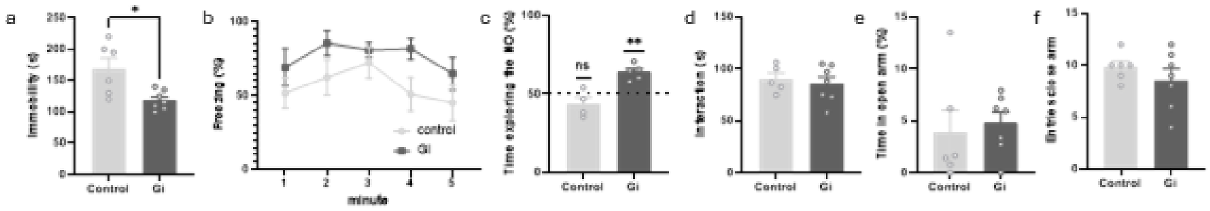


Supplementary figure 2: Order-effect on engram inhibition across extended behavioural measures. a Immobility time in the forced swim test, t-test [t(11)=3.030, p=0.0114], n=6-7. b Freezing levels by minute during fear recall, two-way repeated measures ANOVA [Interaction: F(4, 40) = 0.5087, p=07296; Time: F(4, 40) = 2.528, p=0.0555; AAV: F(1, 10) = 3.544, p=0.0891], n=6-7. c Percentage of time exploring the novel object in the novel object recognition, one sample t-test [Control: t(3)=1.511, p=0.2281; Gi: t(4)=6.259, p=0.0033], n=4-5. d Social interaction with an unknown mice, t-test [t(10)=0.4808, p=0.6410], n=5-7. e Percentage of time in the open arm of the elevated plus maze, t-test [t(11)=0.3718, p=0.7171], n=6-7. f Number of entries in the close arms if the elevated plus maze, t-test [t(11)=1.000, p=0.3386], n=6-7. Data are presented as mean ± SEM.
